# Unacylated Ghrelin Counteracts Contractile and Mitochondrial Dysfunction in Cancer Cachexia

**DOI:** 10.1101/2025.04.29.649515

**Authors:** Bumsoo Ahn, Jonathan Wanagat, Caroline Cleary, Hannah C. Ainsworth, Eunyoung Kim, Hyunyoung Kim

**Author notes:** Corresponding Author: Bumsoo Ahn, PhD, Wake Forest University School of Medicine, Department of Internal Medicine, Medical Center Blvd., Winston-Salem, NC 27157.

## Abstract

**Background:** Cancer cachexia is a complex metabolic syndrome that severely impacts patient mobility, treatment strategies, and quality of life. However, no treatments are available to mitigate the debilitating consequences of cancer cachexia. Unacylated ghrelin (UnAG), the main circulating form of ghrelin, enhances muscle growth and mitochondrial function in various diseases, but its effects in cancer cachexia remain to be tested.

**Methods:** Male C57Bl6/N mice were assigned to one of three treatment groups: non-tumor-bearing (NTB), tumor-bearing (TB), or tumor-bearing treated with unacylated ghrelin (TB+UnAG). Over four weeks, we monitored body weight, food intake, and tumor size. We assessed muscle mass, contractility, mitochondrial oxygen consumption rate (OCR), and reactive oxygen species (ROS) production. Proteomic analysis was performed to elucidate the downstream effects of UnAG. Cell culture assays were performed to measure the in vitro effects of cancer cell-secreted factors and UnAG on myoblasts.

**Results:** Gastrocnemius and quadriceps muscle masses were reduced by 20-30% in TB mice compared to NTB controls; however, UnAG treatment prevented approximately 50% of this loss. Beyond muscle mass, UnAG enhanced the isometric maximum specific force of the extensor digitorum longus by 70% in TB mice. This improvement in muscle quality was associated with preferential upregulation of myosin heavy chain expression in TB+UnAG mice. UnAG also increased mitochondrial OCR while reducing ROS production. Mitochondrial DNA (mtDNA) copy number, which was reduced in TB mice, was restored by UnAG, while the reduced mtDNA mutation frequency in TB mice was maintained with treatment, indicating improved mtDNA integrity. Consistent with enhanced mitochondrial function, treadmill running time was significantly increased in TB+UnAG mice. Proteomic analysis revealed that UnAG downregulated proteins associated with proteolysis, while normalizing antioxidant enzyme thioredoxin and proteins involved in calcium handling. Cancer cell-conditioned medium reduced myotube width in vitro, but UnAG treatment preserved myotube structure..

**Conclusion:** UnAG protects against cancer cachexia by targeting multiple risk factors, including myosin heavy chain expression, mitochondrial bioenergetics, and modulation of protein degradation pathways.

## Introduction

Cancer cachexia is a complex metabolic syndrome characterized by skeletal muscle wasting in the presence of underlying malignancy. Fearon et al. [1] describe it as a multifactorial condition involving persistent skeletal muscle loss with or without loss of fat mass, which cannot be reversed by standard nutritional support. Between 50% and 80% of patients with advanced cancer develop cachexia, with highest incidence reported in patients with lung and gastrointestinal cancer [2]. Cancer cachexia worsens the adverse effects of cancer therapy and diminishes therapy efficacy, thus limiting treatment options for patients with cancer. Cachexia is also associated with poor quality of life and worse overall survival [3]. Metabolic abnormalities, especially impaired mitochondrial bioenergetics and excess free radical generation, are among the key etiological factors contributing to muscle wasting, loss of strength, and fatigue [4]. Despite the high prevalence and consequences of cancer cachexia, no therapeutic options exist to date, highlighting the need for novel interventions targeting the metabolic imbalances and mitochondrial dysfunction driving cancer cachexia.

Ghrelin is a peptide hormone consisting of 28 amino acids primarily produced by X/A-like cells in the stomach. A small subset of ghrelin undergoes acylation, upon which acylated ghrelin binds to the growth hormone secretagogue receptor 1a (GHSR1a) in the pituitary [5]. This interaction stimulates appetite and promotes the release of insulin-like growth factor-1 (IGF-1), an anabolic peptide [5]. Because of GHSR1a’s orexigenic and anabolic effects, GHSR1a receptor agonists (i.e., anamorelin and ONO-7643) were tested in several clinical trials to evaluate their ability to mitigate muscle wasting and loss of strength in patients with cancer [6-8]. In randomized, double-blinded, Phase 3 clinical trials of patients with non-small cell lung cancer, 3- and 6-month treatments with GHSR1a receptor agonists, both anamorelin and ONO-7643, increased lean body mass but failed to increase strength [6-8]. The modest gains in lean body mass and the absence of strength improvements observed in these trials have limited the clinical utility of GHSR1a receptor agonists for patients with cancer. Notably, wildtype mice treated with GHSR1a agonist (HM01) increased fat mass more than lean mass, which resulted in loss of muscle mass and strength [9]. Collectively, existing research reports limitations of applying GHSR1a agonists in ameliorating the symptoms of cancer cachexia patients and mouse models.

Unacylated ghrelin (UnAG) directly affects skeletal muscle and mitochondria without binding to the GHSR1a receptors or causing subsequent increases in appetite or fat mass [10-15]. For example, UnAG incubated with cultured myotubes increases differentiation and fusion of myoblasts via activation of protein synthesis pathways (i.e., mTORC2) [16]. UnAG treatment in vivo induces hypertrophy following acute pathological conditions via downregulation of proteolytic pathways, including FoxO3a and E3 ligases (i.e., MuRF1 and atrogin1) [15]. Furthermore, UnAG improves mitochondrial electron transport system (ETS) activities in in vivo experimental models where mitochondrial defects play a key role in the loss of muscle mass and strength, including diabetes, high fat diet, and chronic kidney disease [13]. Note that the biological effects of UnAG may be mediated through receptor-mediated activation of UnAG downstream pathways. Previous studies suggest the existence of membrane-bound binding sites for UnAG in muscle cells [16, 17], although specific receptors responsible has yet to be identified. Taken together, previous research demonstrated UnAG as an intervention to prevent muscle loss and mitochondrial dysfunction induced by acute and chronic pathological conditions, but there is little information about UnAG’s impact on contractile properties and muscle strength in cancer cachexia, as well as the mechanisms underlying UnAG’s protective impacts.

Our study investigates the potential of UnAG to mitigate wasting, mitochondrial dysfunction, and contractile impairment in tumor-bearing mice. By administering exogenous UnAG to elevate plasma levels, we assessed its effects on muscle physiology, including muscle mass, mitochondrial function, and contractile properties. Additionally, mechanistic experiments using cultured myoblasts were employed to explore the influence of tumor cell-derived circulating factors on muscle growth and the direct impact of UnAG. These findings provide novel insights into the role of UnAG in preserving muscle function in the context of cancer cachexia.

## Materials and methods

### Animals

All animal procedures and experiments were approved by the Institutional Animal Care and Use Committee at the Wake Forest University School of Medicine (Protocol A22-011). We used 4-5-month-old male C57BL6/N mice. The mice were housed under a standard light:dark (12h:12h) cycle and provided ad libitum access to standard chow and water. Mice were assigned to three groups, non-tumor-bearing (NTB), tumor-bearing (TB), and tumor-bearing with unacylated ghrelin treatment (TB+UnAG). Mice in the TB+UnAG group received UnAG in their drinking water at a dose of 100 µg/kg body weight/day, a concentration previously shown to be effective in mitigating skeletal muscle pathology in conditions such as sarcopenia [18] and chronic kidney disease [13]. UnAG-dissolved water was provided for seven days following the Lewis Lung Carcinoma (LLC) inoculation until the endpoint.

### Cancer cell culture

Lewis Lung Carcinoma (LLC, CRL-1642) cells were purchased from the ATCC (Rockville, Maryland, USA) and cultured under standard conditions at 37 °C in a humidified atmosphere containing 5% CO_2_. Cells were grown in DMEM medium (Thermo Fisher Scientific, USA) supplemented with 10% fetal bovine serum, 2 mM glutamine, 100 UI/ml penicillin+0.1 mg/ml streptomycin (Thermo Fisher Scientific, USA). LLC cells at passages 5-10 were suspended in 100 uL of PBS for injection.

### Tumor cell inoculation

C57BL/6N mice, aged 4-5 months, were inoculated subcutaneously with 1×10^6^ LLC cells in 100 μL of PBS, administered at the dorsal midline under the influence of anesthesia. The mice in the NTB group were injected with 100 uL of PBS. Tumors became palpable within one week after injection. The treatment with UnAG (100 ug/kg BW) started one week after the inoculation. In vivo behavioral assessments were conducted two-three days preceding the scheduled euthanasia. We dissected peripheral tissues, including gastrocnemius, quadriceps, soleus, tibialis anterior, *e*xtensor digitorum longus (EDL), heart, and subcutaneous and epididymal adipose tissue. Tumor size and body and tissue weights were recorded.

### Graded exercise tolerance test on treadmill

Exercise tolerance was tested using a graded exercise protocol on a treadmill as previously published [19]. After a 5-minute habituation period, treadmill speed (m/min) and/or duration (min) were increased as follows: 5 m/min (10 min), 7 m/min (10 min), 10 m/min (50 min), and 12 m/min (50 min). A brush located at the end of treadmill was effective in engaging animals on the treadmill. The mice ran until exhaustion, which was determined by failure to engage the treadmill for five seconds in the presence of the brush. We previously found this method of motivating animals to be an effective alternative to electric shocks [19].

### In vitro contractile properties

In vitro contractile function was measured using isolated EDL muscle as previously published [20]. Mice were euthanized using gaseous carbon dioxide, and one EDL muscle was immediately excised and prepared for functional assays in a bicarbonate-buffered solution gassed with a mixture of 95% O_2_ and 5% CO_2_ at room temperature. The EDL muscle was placed in an organ bath containing bicarbonate-buffered solution at room temperature, and the proximal and distal tendons of the EDL were hung on a hook and a force transducer. Muscles were allowed 10 min of thermal equilibration at 32°C and then measurements of force-frequency were initiated. In all electrical stimulations, a supramaximal current (600–800 mA) of 0.25 ms pulse duration was delivered through a stimulator (Aurora Scientific Inc., 701C). All data were recorded and analyzed using commercial software (DMC and DMA, Aurora Scientific, Aurora, Canada). Specific force (N/cm^2^) was calculated by force and estimated fiber cross-sectional area (CSA in cm^2^), based on the fiber length/muscle length ratio of the EDL [21].

### Myofiber permeabilization

Preparation for myofiber permeabilization was performed as previously described [22]. Briefly, a small piece (∼3-5 mg) of red gastrocnemius muscle was carefully dissected, and fibers were separated in ice-cold BIOPS containing 10 mM Ca-EGTA, 0.1 µM free calcium, 20 mM imidazole, 20 mM taurine, 50 mM K-MES, 0.5 mM DTT, 6.56 mM MgCl_2_, 5.77 mM ATP, and 15 mM phosphocreatine at pH 7.1. The muscle bundle was permeabilized in saponin solution (30 µg/mL) for 30 mins, followed by three 5-min washes in ice-cold MiR05 wash buffer containing 0.5 mM EGTA, 3 mM MgCl_2_•6H_2_O, 20 mM taurine, 10 mM KH_2_PO_4_, 20 mM HEPES, 1 g/L BSA, 60 mM potassium-lactobionate, and 110 mM sucrose at pH 7.1.

### Simultaneous measures of mitochondrial respiration and reactive oxygen species production

Oxygen consumption rate (OCR) and reactive oxygen species (ROS) production were simultaneously determined using the Oxygraph-2k (O2k, OROBOROS Instruments, Innsbruck, Austria) as previously described [23]. OCR was determined using an oxygen sensor, and rates of hydrogen peroxide generation were determined using the O2k-Fluo LED2-Module Fluorescence-Sensor Green. Measurements were performed on permeabilized fibers in MiR05 media at 37 °C containing 10 µM Amplex® UltraRed (Molecular Probes, Eugene, OR), 1 U/mL horseradish peroxidase (HRP), SOD (5 U/mL), and 25 µM blebbistatin. HRP catalyzes the reaction between hydrogen peroxide and Amplex UltraRed to produce the fluorescent resorufin (excitation: 565 nm; emission: 600 nM). The fluorescent signal was converted to nanomolar H_2_O_2_ via a standard curve established on each day of experiments. Background resorufin production was subtracted from each measurement. Rates of respiration and ROS production were determined using sequential additions of substrates and inhibitors as follows: glutamate (10 mM), malate (2 mM), ADP (5 mM), succinate (10 mM), rotenone (1 µM), and Antimycin A (1 µM). All respiration measurements were normalized to Antimycin A to account for non-mitochondrial oxygen consumption. Data for both OCR and rates of ROS production were normalized by milligrams of muscle bundle wet weights.

### mtDNA copy number and deletion mutation frequencies

MtDNA copy number and deletion mutation frequency were measured using previously validated digital PCR approaches [24]. Gastrocnemius muscles were powered with a mortar and pestle under liquid nitrogen. We extracted total DNA from frozen muscle powder using a DNA extraction kit (GenFind V3, Beckman Coulter, Lifesciences, CA, USA) and eluted in 10 mM Tris-EDTA buffer, pH 8. The quality and quantity of total DNA were assessed using spectrophotometry at A230, A260, and A280 (Nanodrop 2000 Spectrophotometer, ThermoScientific, MA, USA) and fluorometry (Qubit 2.0 Fluorometer, ThermoScientific, MA, USA). DNA integrity was examined by gel electrophoresis or Tapestation (TapeStation 4200, Agilent, CA, USA). We used a 5-prime nuclease cleavage assay and droplet-digital PCR (ddPCR) to quantify copy numbers for nuclear DNA, total mtDNA, and mtDNA deletion mutations with specific primer/probe sets for each as previously described [24]. We diluted the samples to fall within the manufacturer’s recommended range (20 to 2,000 target copies per microliter). Digital PCR cycling conditions included polymerase activation at 95°C for ten minutes, 40 cycles of denaturation at 94°C for 30 seconds, and annealing/extension at 60°C for two minutes. We determined reaction thresholds and target copy number per microliter values using QuantaSoft Software (Bio-Rad,CA, USA). The same cycling conditions were used for direct quantitation of the mtDNA major arc deletions by ddPCR but extended to 60 cycles. Researchers performing the mtDNA assays were blinded to sample characteristics.

### Myofibrillar gels

To solubilize myosin and actin proteins, small pieces of gastrocnemius samples (3-5 mg) were homogenized in a high-salt lysis buffer, previously shown to be optimal for myosin and sarcomeric actin extraction and solubility [25]. The buffer consisted of 300 mM NaCl, 0.1 M NaH_2_PO4, 0.05 M Na_2_HPO_4_, 0.01 M Na_4_P_2_O_7_, 1 mM MgCl_2_, 10 mM EDTA, 1 mM DTT (pH 6.5), and complete protease inhibitor cocktail (Sigma-Aldrich). Tissue lysates were subsequently centrifuged at 16,000 g for 3 min at 4 °C, and the supernatant was collected. The samples were mixed with Laemmli buffer (Bio-Rad) and boiled before loading. To determine MyHC abundance, we loaded 0.6 μg protein/lane (which was within the linear range determined in our laboratory) into a 10% polyacrylamide gel (Criterion precast gels; Bio-Rad) and ran the gel electrophoresis at 200 V for 50 min at 4 °C. The gel was stained with Coomassie blue for 3 h (Thermo Fisher Scientific, Waltham, MA, USA) and washed with ddH_2_O overnight. We quantified the optical density corresponding to MyHC and actin using an Odyssey Infrared Imaging system (LI-COR, Lincoln, NE, USA).

### qRT PCR

Total RNA was extracted from myoblasts using TRIzol reagent (Invitrogen, Carlsbad, CA, USA). RNA purity and yield were determined by measuring the absorbance at 260 and 280 nm using a Nano Drop. The first-strand cDNA was synthesized from 1 µg total RNA using iScript™ cDNA synthesis kit (Bio-Rad, Hercules, CA, USA) and 5 ng cDNA samples were amplified using TaqMan SYBR master mix (Thermofisher, USA). Primers are listed in the Supplemental Table. Quantitative Real-time PCR (qRT-PCR) was performed using the QuantStudio 5 real-time PCR instrument (Thermo Fisher scientific, USA). The qRT-PCR data were analyzed using QuantStudio™ Design & Analysis software. The threshold cycle (Ct) values were determined, and the relative gene expression levels were calculated using the ΔΔCt method, normalizing to the expression of beta-actin in this experiment.

### Preparation of cancer cell-conditioned medium

To prepare cancer cell-conditioned medium (CM), LLC cells were seeded and cultured in DMEM with 10% FBS as previously described [26]. Once LLC cells reached 70-80% confluence in a growth medium, the cells were washed with PBS and cultured in serum-free DMEM for 24 h. The medium was collected and centrifuged at 3,000 rpm for 10 min, and the supernatant was collected in a fresh tube to be either used immediately or stored at −80 °C for future use. CM was prepared from an equal number of cancer cells for each cell line. CM was reconstituted with 2% horse serum and 1 µg/ml insulin before treating myotubes.

### C2C12 cell culture, differentiation, and treatment

The C2C12 mouse myoblasts were obtained from the ATCC (Rockville, Maryland, USA). Cells were cultured in dishes and maintained in high glucose Dulbecco’s modified Eagle’s medium (DMEM) supplemented with 10% fetal bovine serum, 2 mM glutamine, and 100 UI/ml penicillin+0.1 mg/ml streptomycin at 37 °C in a humidified atmosphere of 5% CO_2_. The culture medium was refreshed every 48 hours. For differentiation, C2C12 cells were plated at a density of 100 cells/mm^2^. Confluence was reached three days after seeding, and myogenic differentiation was induced by replacing the growth medium with differentiation medium, DMEM containing 2% horse serum. The medium was replaced every 24 hours.

### Myotube measures

To analyze myotube width of differentiation medium (DM)-treated, differentiation medium-conditioned medium (DMCM)-treated, DMCM+UnAG-treated myotubes, 10 images were obtained for each group. 25∼50 myotubes were analyzed for each image using image J software. The width of myotubes was calculated as the average from three measures per myotube.

### Global proteomics

Snap frozen gastrocnemius samples were analyzed on an LC-MS/MS system consisting of an Orbitrap Eclipse Mass Spectrometer (Thermo Scientific, Waltham, MA, USA) and a Vanquish Neo nano-UPLC system (Thermo Scientific, Waltham, MA, USA). Peptides were separated on a DNV PepMap Neo (1500 bar, 75 μm x 500 mm) column for 120 min employing linear gradient elution consisting of water (A) and 80% acetonitrile (B) both of which contained 0.1% formic acid. Data was acquired by top speed data dependent mode where maximum MS/MS scans were acquired per cycle of 3 seconds between adjacent survey spectra. Dynamic exclusion option was enabled for which the duration was set to 120 seconds. To identify proteins, spectra were searched against the UniProt mouse protein FASTA database (17,082 annotated entries, Oct 2021) using the Sequest HT search engine with the Proteome Discoverer v2.5 (Thermo Scientific, Waltham, MA). The search parameters were as follows: FT-trap instrument; parent mass error tolerance, 10 ppm; fragment mass error tolerance, 0.6 Da (monoisotopic); enzyme, trypsin (full); # maximum missed cleavages, 2; variable modifications, +15.995 Da (oxidation) on methionine; static modification (only for soluble part), +57.021 Da (carbamidomethyl) on cysteine. Samples with high degree of missingness, relative to other samples, were dropped from analyses and proteomics analyses were limited to proteins that were detectable across all samples. Protein intensities were normalized and log_2_ transformed for analyses.

### Statistical analysis

The data were analyzed by Prism 10.0 (GraphPad, La Jolla, CA, USA). Pairwise comparisons involved the use of unpaired two-tailed t-tests to compare differences between groups. For multiple groups, ordinary One-Way and Two-Way ANOVAs were performed with Tukey post hoc tests as indicated in the figure legends. For each outcome, statistical significances were declared at p <0.05. Data are presented as mean ± Standard Error of Mean (SEM). For proteomics analyses, comparisons were limited to TB vs TB+UnAG samples using a two-sample t-test. Statistical significance was determined using the Benjamini-Hochberg False Discovery Rate (BH-FDR), but suggestive statistical support was declared at p <0.05.

## Results

### UnAG preserves muscle mass in cancer cachexia without affecting tumor growth or food intake

Tumor growth and food consumption were measured following exogenous UnAG administration in TB mice, with no significant differences observed between the TB and TB+UnAG groups (Fig. 1a, b). Tumor free body weights were reduced by 12-15% in both groups (Fig. 1c). Notably, gastrocnemius and quadriceps muscle masses were reduced by 20-30% in TB mice; however, UnAG treatment preserved the muscle mass by 50% (Fig. 1d, f). This protective effect remained significant after normalization by tibial length (Fig. 1e, g). In contrast, fat mass was reduced by ∼70% in TB mice, and UnAG had minimal impact on this loss (Fig. S1). Collectively, UnAG did not alter tumor volume or food intake of TB mice but maintained muscle mass.

**Figure 1.**
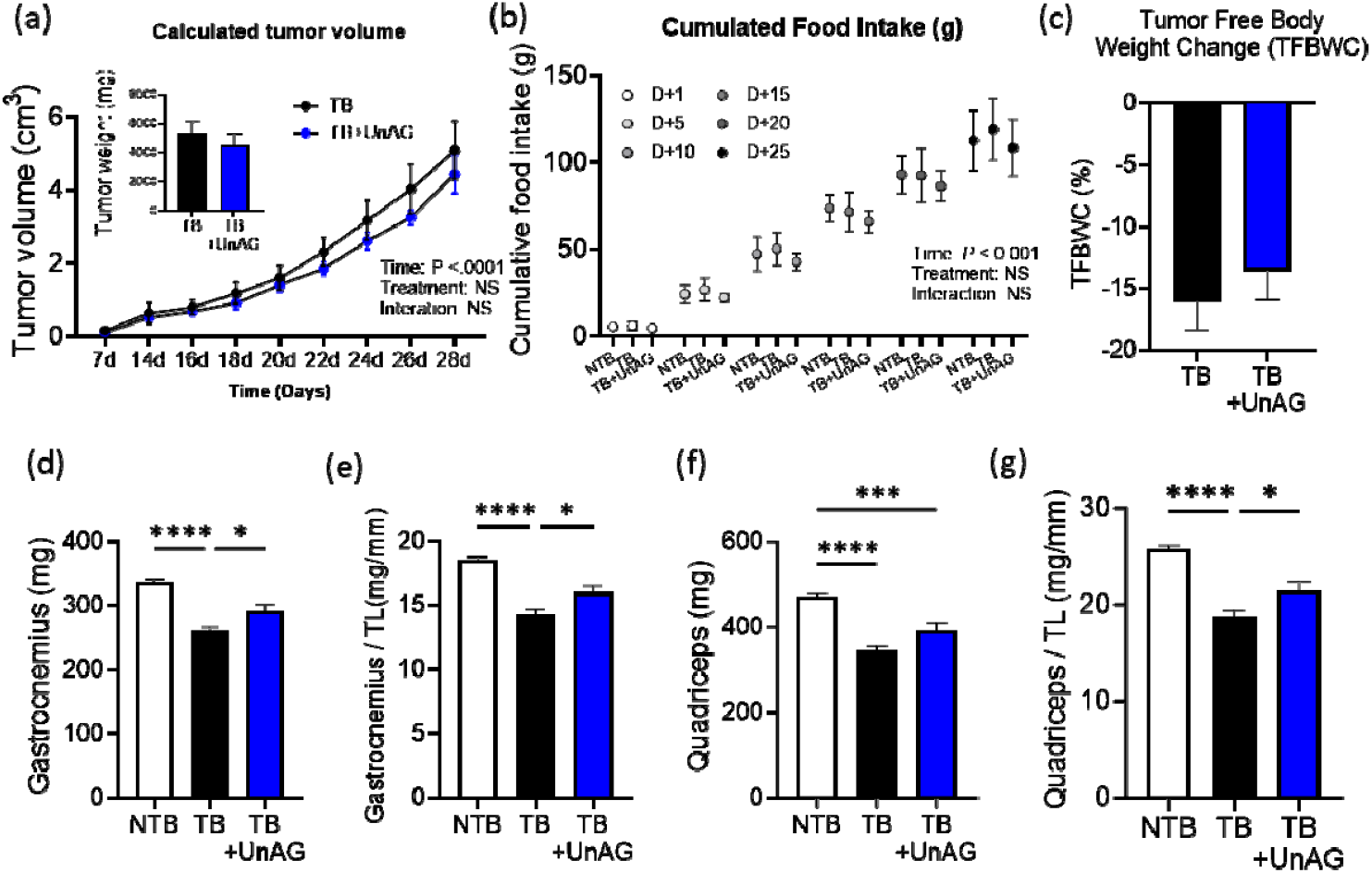
Effects of UnAG on tumor volume, food consumption, and lower limb muscle mass of tumor bearing mice. (a) Tumor volume (cm^2^) in the tumor bearing mice did not differ by UnAG treatment. Inset indicates tumor weights at the end point. n=5-6. One-way ANOVA. (b) Cumulative food intake at different timepoints after tumor cell inoculation. n=4-8. One-way ANOVA. (c) Tumor free body weight at 28 days post tumor cell inoculation. n=9-13. (d) Gastrocnemius muscle mass from both legs in mg and (e) gastrocnemius mass relative to tibial length. n=10-15. (f) Quadriceps muscle mass from both legs in mg and (g) quadriceps mass relative to tibial length. n = 10-15. One-way ANOVA followed by Tukey post hoc tests. \**P*<.05. \*\**P*<.01. \*\*\**P*<.001. \*\*\*\**P*<.0001. Bars represent means ±SEM. Abbreviations: TB, tumor bearing; NTB, non-tumor bearing; UnAG, unacylated ghrelin; TL, tibial length.

### UnAG improves contractile dysfunction of skeletal muscle in TB mice

Another key component of cancer cachexia, in addition to wasting, is loss of strength. Isometric contractile properties of the extensor digitorum longus (EDL) muscle were assessed by generating a force–frequency curve (Fig. 2a). The maximum isometric specific force, measured as force per CSA, was reduced by 30% in the TB mice, but UnAG treatment rescued 70% of the force deficit in the TB mice (Fig. 2b). In the analysis of twitch characteristics, we found that peak twitch force and time to half relaxation (1/2RT) did not differ by tumor or UnAG (Fig. 2c, d). Time to peak twitch was longer for TB mice compared to NTB, but UnAG treatment shortened the time to peak twitch similar to NTB level (Fig. 2e), suggesting the improvement in calcium handling mechanisms by UnAG. Our results demonstrate that UnAG improves contractile function in TB mice.

**Figure 2.**
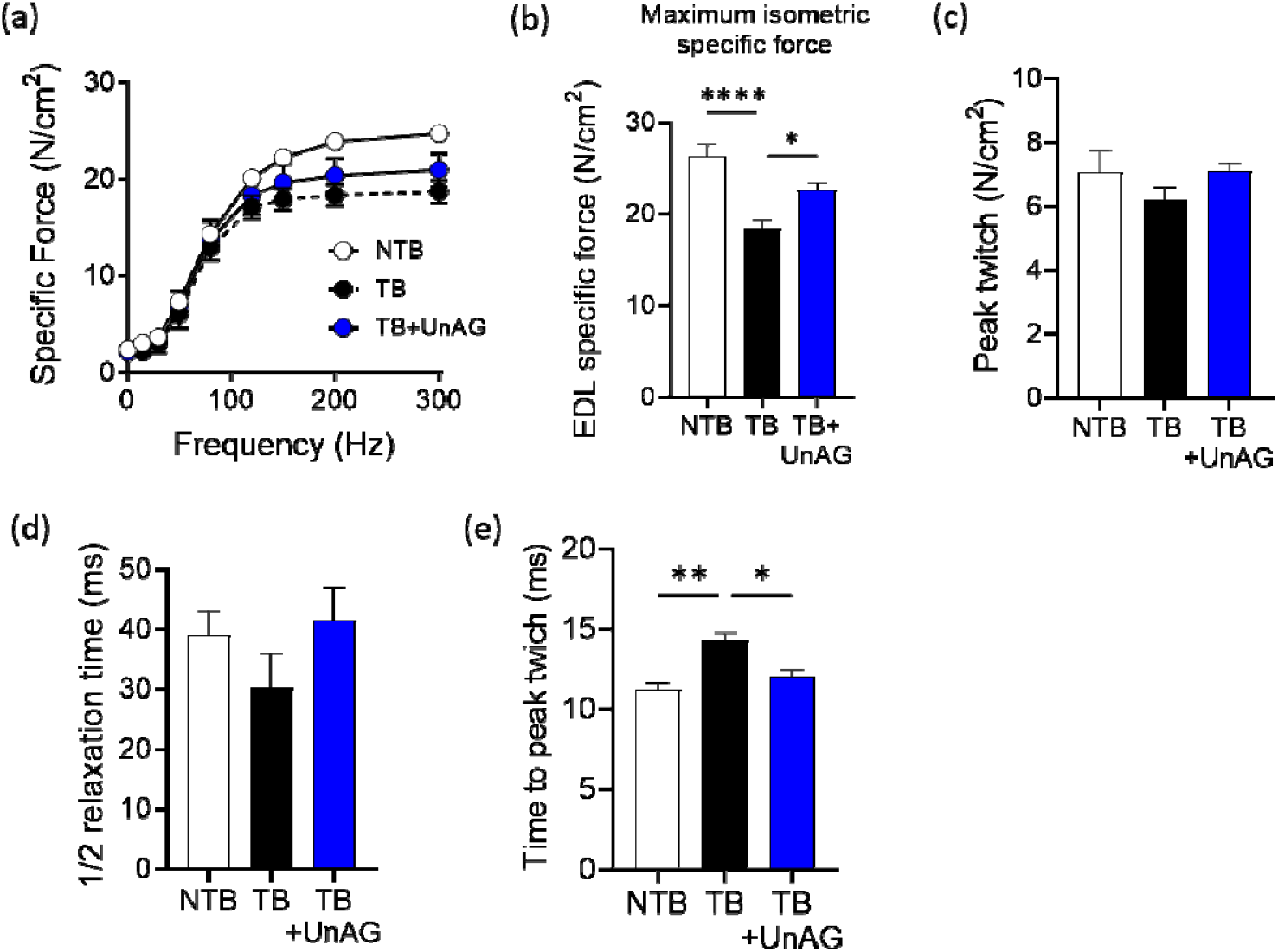
In vitro contractile properties of isolated skeletal muscle in tumor bearing mice. Force–frequency relationship was assessed in isolated extensor digitorum longus (EDL) muscles to evaluate contractile responsiveness. (b) Maximum isometric specific force. (c) Twitch tension (N/cm2), (d) one half relaxation time (1/2 RT), and (e) time to peak twitch. n=5-6. One-way ANOVA followed by Tukey post hoc tests. \**P*<.05. \*\**P*<.01. \*\*\*\**P*<.0001. Bars represent means ±SEM. Abbreviations: TB, tumor bearing; NTB, non-tumor bearing, UnAG, unacylated ghrelin.

### UnAG increases MyHC expression of TB mice

To determine whether the impairment in maximum specific force is linked to the alterations in myofibrillar protein expression, we quantified key sarcomeric proteins MyHC and actin using the whole tissue homogenates of gastrocnemius muscle. We used a high-salt buffer that demonstrated increased solubility of myofibrillar proteins [27]. We found that MyHC levels decreased by ∼30% in the TB mice compared to NTB controls, whereas actin levels remained unchanged (Fig. 3a, b). This preferential loss of MyHC has been reported in the diaphragm of C-26 TB mice [25]. Notably, UnAG treatment prevented MyHC loss in TB mice, suggesting that its protective effects may be linked to enhanced myogenesis or reduced protein degradation of MyHC in skeletal muscle, as shown in other pathological conditions [12, 14, 15].

**Figure 3.**
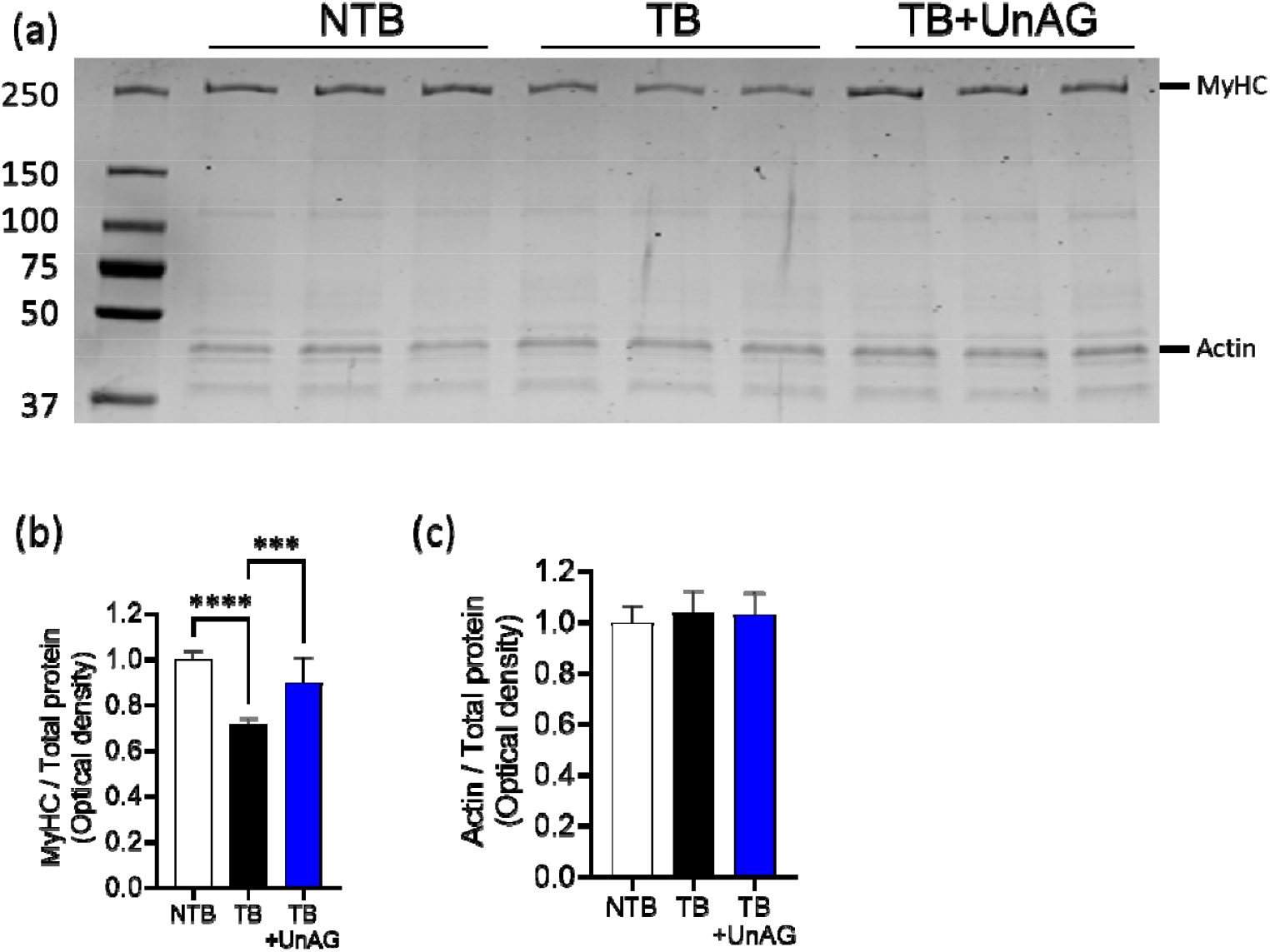
Myosin heavy chain (MyHC) expression in the gastrocnemius muscle of NTB, TB, and TB-UnAG groups. (a) Representative gel showing MyHC and actin expression. (b) Densitometric quantification of MyHC expression, normalized to total protein within the solubilized fraction. (c) Densitometric quantification of actin expression, normalized to total protein within the solubilized fraction. n=6. One-way ANOVA followed by Tukey post hoc tests. \*\*\**P*<.001. \*\*\*\**P*<.0001. Bars represent means ±SEM. Abbreviations: MyHC, myosin heavy chain.

### UnAG enhances mitochondrial respiration while downregulating ROS production in TB mice

Defects in mitochondrial bioenergetics and excess ROS are well documented in TB mice and in patients with cancer [28, 29]. Given the known effects of UnAG on the mitochondrial respiration and ROS generation [10, 11, 13], we assessed mitochondrial OCR and ROS production rates using permeabilized gastrocnemius myofibers. Representative traces in Fig 4a illustrate mitochondrial responses to sequential substrate and inhibitor additions, with all experiments completed within a 50-minute timeframe to preserve myofiber integrity. In TB mice, ADP-stimulated complex I- and II-driven oxidative phosphorylation (i.e., OxPhos capacity) was reduced by ∼40% compared to NTB mice but was normalized by UnAG treatment. Similarly, complex II-driven respirations in TB mice were reduced by ∼50% compared to NTB mice but were restored by UnAG (Fig. 4b). Mitochondrial ROS production, which was elevated across all substrate and inhibitor conditions in TB mice, was also normalized following UnAG treatment (Fig. 4c). Furthermore, mitochondrial DNA (mtDNA) copy number decreased by 33% in TB mice compared to NTB controls but was restored by UnAG (Fig. 4d). Interestingly, mtDNA deletion frequency was lower in both TB and TB+UnAG groups compared to NTB mice (Fig. 4e). Finally, exercise tolerance, assessed via a graded exercise test, was reduced by 33% in TB mice but was fully restored by UnAG treatment (Fig. 4f). These findings highlight the potential of UnAG to mitigate mitochondrial dysfunction and improve oxidative metabolism in TB mice, ultimately supporting muscle function and exercise capacity.

**Figure 4.**
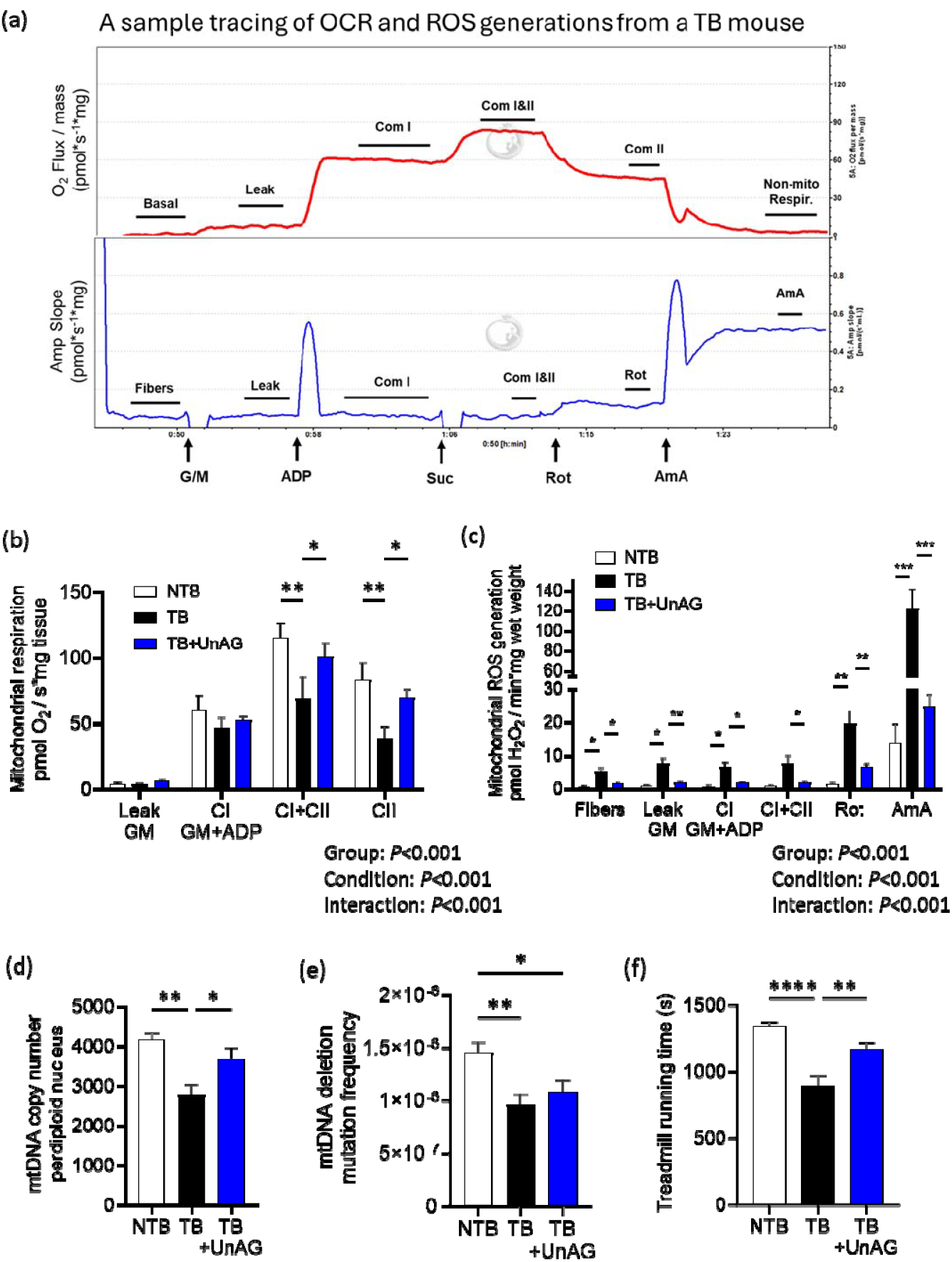
Mitochondrial OCR and ROS production of permeabilized myofibers from tumor bearing mice. (a) A sample tracing of OCR and ROS generation rates. Average values of OCR (b) and ROS (c) generation rates in response to substrates and inhibitors of individual complexes. n=6-10. Two-Way ANOVA followed by Tukey post hoc. (d) mitochondrial DNA copy numbers and (e) mitochondrial DNA deletion mutation frequency. N=7. (f) Treadmill running time in response to graded exercise test. n=7-13. One-way ANOVA followed by Tukey post hoc tests. \**P*<.05. \*\**P*<.01. ***P<.001. \*\*\*\**P*<.0001. Bars represent means ±SEM. Abbreviations: OCR, oxygen consumption rate; ROS, reactive oxygen species; G/M, glutamate+malate; Rot, rotenone; AmA, antimycin A; CI, complex I; CII, complex II; CIV, complex IV; mtDNA, mitochondrial DNA.

### Proteome analyses reveal key proteins and pathways modulated by UnAG

Proteomic quantification identified 2,572 proteins across all samples (NTB, TB, TB+UnAG) passing quality control metrics, with only one NTB sample removed due to a high percentage of protein missingness. While no individual proteins met significance after BH-FDR adjustment, statistical analysis identified proteins modulated (with suggestive evidence at p <0.05) by UnAG in TB mice, relating to key regulators of muscle size and contractile function, including protein degradation, antioxidant defense, and excitation-contraction (EC) coupling. Five proteins involved in protein degradation pathways were elevated in TB mice but normalized by UnAG treatment (Fig. 5a). Additionally, UnAG reduced the antioxidant thioredoxin in TB mice (Fig. 5b). Since oxidative stress increases antioxidant expression [23, 30], this suggests that UnAG mitigates oxidative stress, aligning with the mitochondrial ROS data in Fig. 4c. UnAG also restored the expression of ryanodine receptor 1 (RyR1), a key regulator of calcium release, to NTB levels (Fig. 5c). Furthermore, UnAG upregulated fast myosin-binding protein-C (fMBP-C), which influences calcium sensitivity in muscle cells [31], bringing its expression to NTB levels in TB mice (Fig. 5d). Collectively, these findings support the hypothesis that UnAG modulates critical proteins involved in protein homeostasis, antioxidant defense, and EC coupling, contributing to muscle function preservation in TB mice.

**Figure 5.**
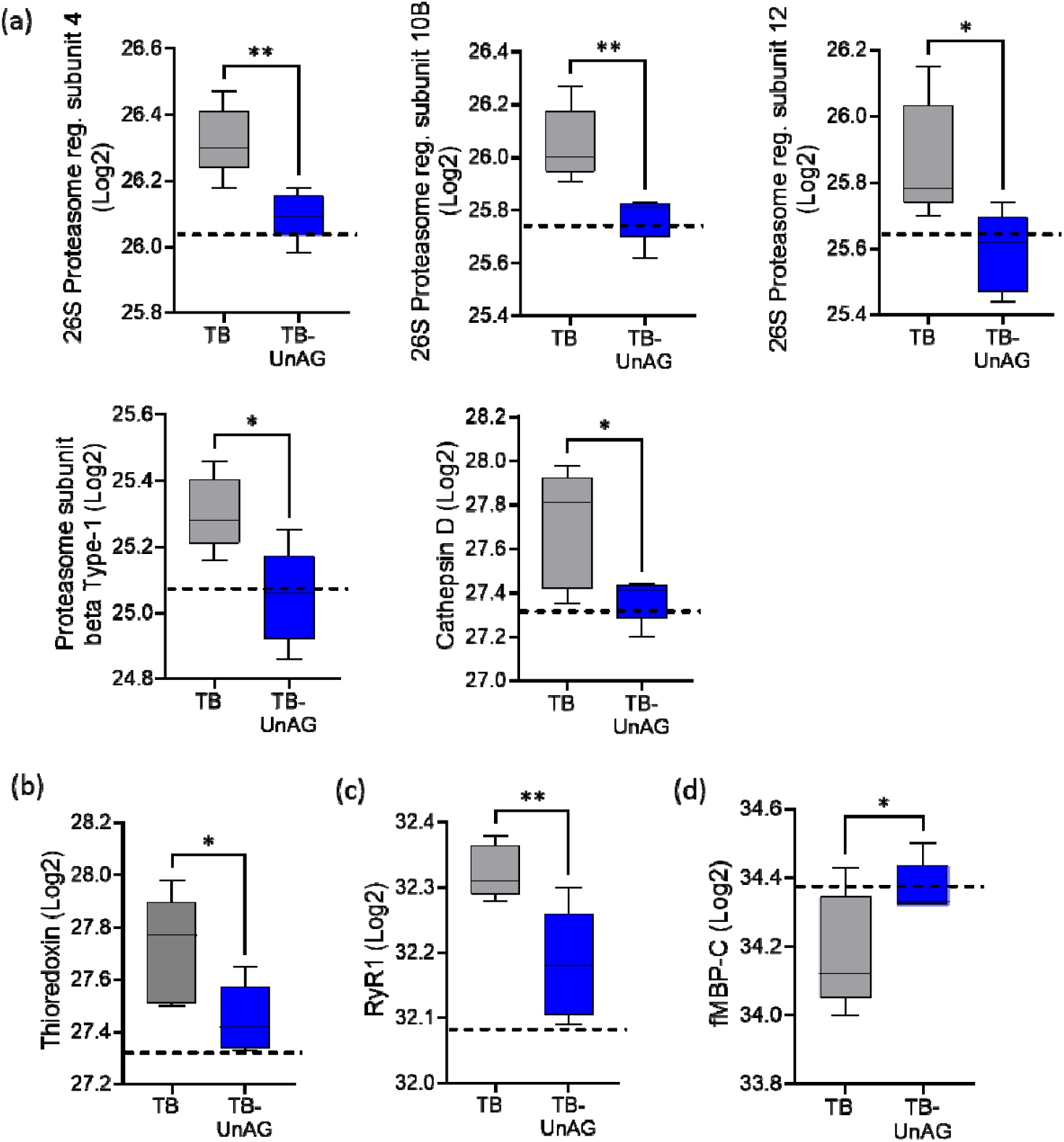
Expression of proteins involved in protein degradation, antioxidant defense system, and EC coupling process in the gastrocnemius. Data are normalized to Log2 scale. n=5 for TB and TB-UnAG. Dotted lines indicate average values from NTB mice (n=4) for reference. (a) Expression of proteins involved in protein degradation, including 26S proteasome regulatory subunits and cathepsin D. (b) Protein expression of antioxidant defense system, thioredoxin. (c) Expression of proteins of RyR1. (d) Protein expression of fMBP-C. Student’s t-tests were used to compare means of each group. While these proteins did not meet BH-FDR significance thresholds, p values are shown as: \**P*<0.05. \*\**P*<0.01. In the box plots, the box represents the interquartile range, encompassing the middle 50% of the data, while the line within the box marks the median. The lines extending from the box (whiskers) show the range of the remaining data. Abbreviations: RyR1, ryanodine receptor1; fMBP-C, fast myosin binding protein-C; EC, excitation-contraction.

### UnAG normalizes reduced protein synthesis and myotube atrophy in cancer cell-conditioned media

To investigate the impact of circulating factors on myogenesis and myotube growth in cancer cachexia, we cultured myoblasts with differentiating media (DM), DM plus cancer cell-conditioned media (DMCM) and DMCM co-incubated with UnAG (DMCM+UnAG). The width of myotubes cultured in DMCM were significantly smaller than those in DM; however, UnAG treatment restored myotube width to control levels (Fig. 6a). Analysis of mRNA expression in protein degradation pathways revealed that UnAG normalized MuRF1 levels observed in DM-cultured myotubes (Fig. 6b), while atrogin1 expression remained unchanged by either CM or UnAG treatment (Fig. 6c, d). UnAG significantly increased eMHC levels in DMCM (Fig. 6e). Additionally, UnAG reduced upregulation of mRNA expressions involved in calcium release and uptake, including SERCA1, phospholamban, calciquestrin, and voltage-dependent calcium channel (Fig. S2). Together, our findings demonstrate that cancer cell-secreted factors contribute to reduced myofiber size and alter the expression of mRNAs involved in calcium handling—effects that are restored by UnAG.

**Figure 6.**
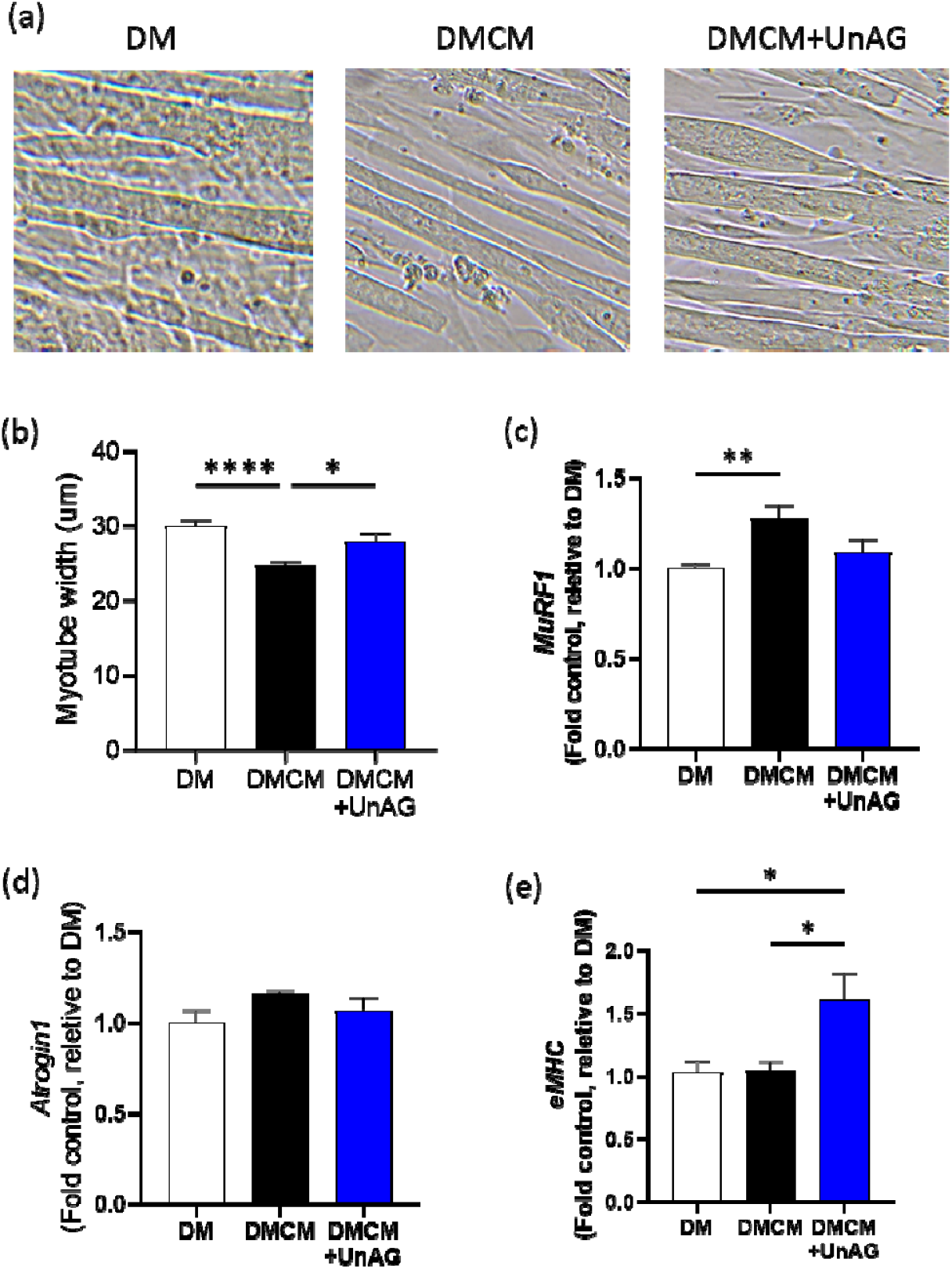
Effects of cancer cell-derived circulating factors and UnAG on cultured myotubes. (a) Brightfield microscopic images of myotubes incubated with differentiating medium (DM), DM plus conditioned medium (DMCM), and DMCM treated with 100 nM UnAG for 24 h. (b) Quantified width of myotubes from DM, DMCM, and DMCM+UnAG. n=10. mRNA expression of genes involved in protein degradation, myogenesis, and denervation, including *MuRF1* (c), *Atrogin1* (d), *eMHC* (e). n=10. One-way ANOVA followed by Tukey post hoc tests. \**P*<0.05. \*\**P*<0.01. \*\*\*\**P*<0.0001. Bars represent means ±SEM. Abbreviations: DM, differentiating medium; DMCM, differentiating medium and conditioned medium; DMCM+UnAG, DMCM treated with unacylated ghrelin.

## Discussion

The aim of this study was to evaluate the therapeutic effects of UnAG on muscle mass, mitochondrial function, and contractile properties in TB mice. Our findings reveal that UnAG preserves muscle mass and prevents contractile dysfunction in TB mice without influencing tumor volume or food intake. These protective effects are associated with increased myofiber size, enhanced MyHC expression, improved mitochondrial bioenergetics, and reduced ROS generation. Our findings highlight UnAG as a promising therapeutic strategy for counteracting cancer cachexia.

Muscle wasting is a prominent characteristic of patients with advanced lung cancer, even in patients with preserved body weights [32], which is also recapitulated in an LLC-derived cancer cachexia model [29]. UnAG treatment attenuated the loss of muscle mass in TB mice without changing tumor size or food consumption. UnAG treatment downregulated the expression of proteins involved in proteolysis, including four proteins in proteasome pathways (Fig. 5a). The 26S proteasome constitutes the central proteolytic machinery in eukaryotic cells that are known to be upregulated with aging and chronic diseases, including cancer cachexia. Several groups showed downregulation of ubiquitin proteasome pathways by UnAG (i.e., FoxO3a and MuRF1) in atrophying muscle elicited by pathological conditions [12, 13, 15]. Our observed downregulation of proteins involved in protein degradation with UnAG treatment is consistent with this previous work and elucidates a mechanism by which UnAG may work to help prevent muscle wasting. UnAG has been shown to activate myogenesis and protein synthesis pathways [16], but proteins involved in these pathways were not altered by UnAG in our TB mice. The effects of UnAG were specific to skeletal muscle as fat mass wasting persisted.

Loss of strength in cachectic cancer patients is correlated with reduced efficacy of chemotherapy and decreased independence. Previous attempts to mitigate cachexia were met with limited success due to a lack of improvement in strength [6-8]. UnAG treatment in TB mice increased the specific force of isolated EDL muscle by 50%, indicating that UnAG may prevent loss of strength in cancer patients. One of the contributors of decreased specific force in cancer cachexia is myofibrillar protein expression. Investigators previously reported mixed results regarding the expression of MyHC in skeletal muscle subject to cancer cachexia [25, 27]. Using C26 TB mice, Cosper et al. reported no evidence of preferential MyHC loss in the gastrocnemius [27], whereas Roberts et al. showed selective loss of MyHC in the diaphragm [25]. Our findings align with the latter, demonstrating a ∼30% reduction in MyHC in the gastrocnemius of LLC TB mice. UnAG treatment fully preserved MyHC expression in TB mice, which correlates with increased specific force and improved inverted grid hanging time. These results are consistent with the role of UnAG in promoting myogenesis and upregulation of MyHC expression in cultured myotubes [16]. The effects of UnAG on protein synthesis and degradation were confirmed in our proteomic analyses. Collectively, our data supports preferential loss of MyHC in the LLC TB mice and highlights the protective effects of UnAG in maintaining MyHC expression.

Excitation-contraction (EC) coupling is a critical mechanism in force modulation, involving motor neuron excitation of the sarcolemma, calcium release via RyR, calcium-activated force generation, and calcium reuptake by SERCA [33]. Our analysis of twitch characteristics indirectly assessed calcium sensitivity (peak twitch), calcium reuptake (half-relaxation time), and calcium release (time to peak tension). We found that time to peak twitch was prolonged in TB mice but restored to normal levels by UnAG, suggesting improved calcium release in EDL muscle [21]. Consistent with this functional data, our proteomic analysis showed that RyR1 expression was upregulated in TB mice and normalized by UnAG. Together, these results indicate that UnAG preserves EC coupling by normalizing calcium handling in TB mice.

Mitochondrial dysfunction and oxidative damage are well-documented in both cancer patients and mouse models of cancer cachexia. In cancer patients who experienced significant loss of body mass over 6–12 months, skeletal muscle exhibited impaired OxPhos capacity and elevated mitochondrial ROS (mtROS) generation [34]. Similarly, our TB mice also exhibited diminished OxPhos capacity and elevated mtROS levels in skeletal muscle. However, UnAG treatment restored mitochondrial respiration and reduced mtROS generation in TB mice to the levels of NTB mice. We anticipate that the beneficial effects of UnAG on the mitochondria may contribute to improved muscle function and mass preservation. This is consistent with previous investigations demonstrating that mitochondria-targeted interventions mitigated myofiber atrophy and weakness in cancer cachexia [35, 36]. Consistently, UnAG treatment also enhanced mitochondrial bioenergetics and reduced mtROS generation in loss of muscle mass and function elicited by chronic kidney diseases [13] and age [9, 10].

Studies on animal models of cancer cachexia have reported a decline in mitochondrial content and mtDNA copy number in muscle [37]. Our results are consistent with these findings, as we observed a 33% reduction in mtDNA copy number in LLC TB mice compared to NTB controls. Notably, UnAG treatment fully prevented or recovered the decreases in mtDNA copy number in TB mice, signifying its positive impact on mtDNA quantity. MtDNA copy number has proven to be an indicator of muscle mass [38], and this aligns with our findings showing the muscle wasting in TB mice and the maintenance of muscle mass in the UnAG treated mice. Investigators have previously demonstrated that mtDNA copy number measured by quantitative PCR weakly correlates with mitochondrial content and mitochondrial function [39]. However, it is possible that the improvement in copy number in the UnAG treated mice may have contributed to our observed improvements in mitochondrial function indirectly through improved mitochondrial quality control mechanisms accompanying the increased mtDNA copy number. Given the declines in mtDNA copy number that occur with cancer cachexia, the ability of UnAG to preserve mtDNA copy number may be highly relevant to mitigating early fatigue and loss of mobility in cancer patients.

Excess free radicals are known to cause oxidative modifications in proteins, lipids, and DNA [19, 22]. Interestingly, our results showed lower mtDNA deletion mutation frequencies in TB mice compared to NTB mice. While unexpected, similar findings have been reported in colorectal cancer cells, where mtDNA mutations are less frequent than in non-tumor tissues [40]. Our results build on this observation by demonstrating that muscle tissue in TB mice also exhibits fewer mtDNA mutations than in non-TB mice. Notably, increased fidelity of the mitochondrial genome was maintained in TB mice treated with UnAG, as UnAG treatment led to increases in mtDNA copy number without increasing mutation frequencies.

The mechanisms by which cancer induces wasting and weakness in TB mice remain unclear. To determine the effects of cancer cell-secreted factors on muscle atrophy, myoblasts were cultured with cancer cell-conditioned medium, which resulted in significant atrophy in myotube width consistent with the literature [26]. Co-incubation with UnAG, however, prevented the atrophy, which was associated with downregulation of MuRF1 expression, one of the key E3 ligase enzymes involved in ubiquitin proteasome pathways. Consistent with our results, others have reported that myotubes incubated with UnAG activates protein synthesis pathways while downregulating protein degradation, which contribute to muscle growth [15, 16].

In summary, our findings demonstrate that UnAG effectively preserves muscle mass, strength, and mitochondrial function in a mouse model of cancer cachexia without affecting tumor volume or food intake. UnAG mitigated myofiber atrophy, restored MyHC expression, and improved mitochondrial bioenergetics while reducing oxidative stress and proteolytic activity. Our results highlight the therapeutic potential of UnAG in counteracting muscle wasting and functional decline associated with cancer cachexia, offering a promising avenue for future interventions aimed at improving patient outcomes.

## Acknowledgement

B.A. was supported by the National Institute of Aging [AG064143]. This work was partially supported by the Wake Forest University Claude D. Pepper Older Americans Independence Cetner [P30AG21332]. We would like to thank pilot funding support from Center for Redox Biology in Medicine (CRBM) and Atrium Health Wake Forest Baptist Comprehensive Cancer Center (AHWFBCCC). The authors wish to acknowledge the support of the Atrium Health Wake Forest Baptist Comprehensive Cancer Center Proteomics and Metabolomics Shared Resource, supported by the National Cancer Institute’s Cancer Center Support Grant award number P30CA012197. JW was supported by the National Institutes of Health (R01AG069924) and the Department of Veterans Affairs (I01RX004521). The content is solely the responsibility of the authors and does not necessarily represent the official views of the National Institutes of Health.

## Data Availability Statement

The data that support the findings of this study are available from the corresponding author upon a reasonable request.

## Conflict of Interest Statement

None declared.

## Supplemental Figures

**Supplemental Figure 1.**
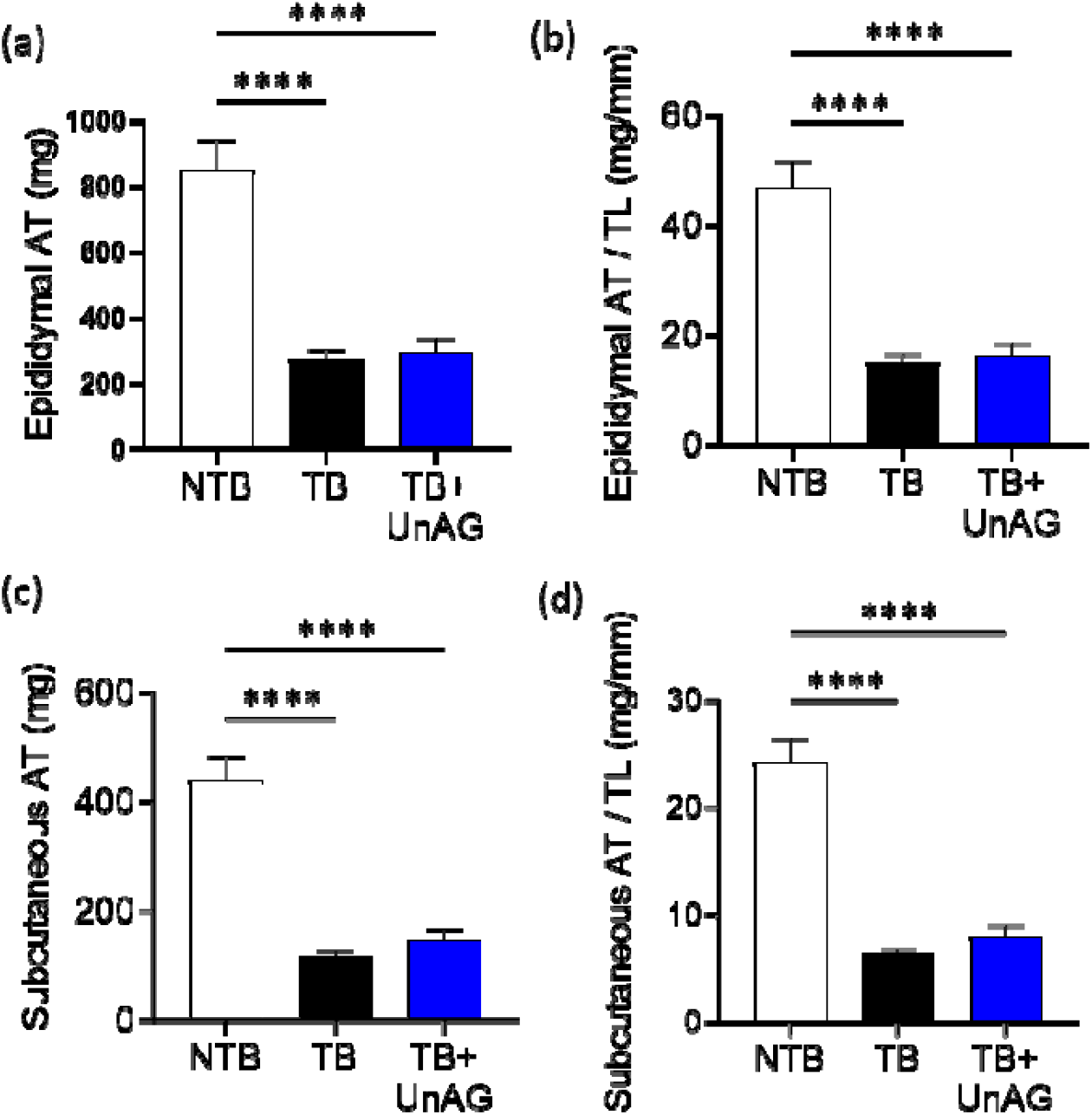
Effects of UnAG on adipose tissue weights. (a) Epididymal adipose tissue weight. (b) Epididymal adipose tissue weights normalized by tibial length. (c) Subcutaneous adipose tissue weight. (d) Subcutaneous adipose tissue weights normalized by tibial length. n=10-15. One-way ANOVA followed by Tukey post hoc tests. ****p<0.0001. Bars represent means ±SEM. Abbreviations: AT, adipose tissue; TL, tibial length.

**Supplemental Figure 2.**
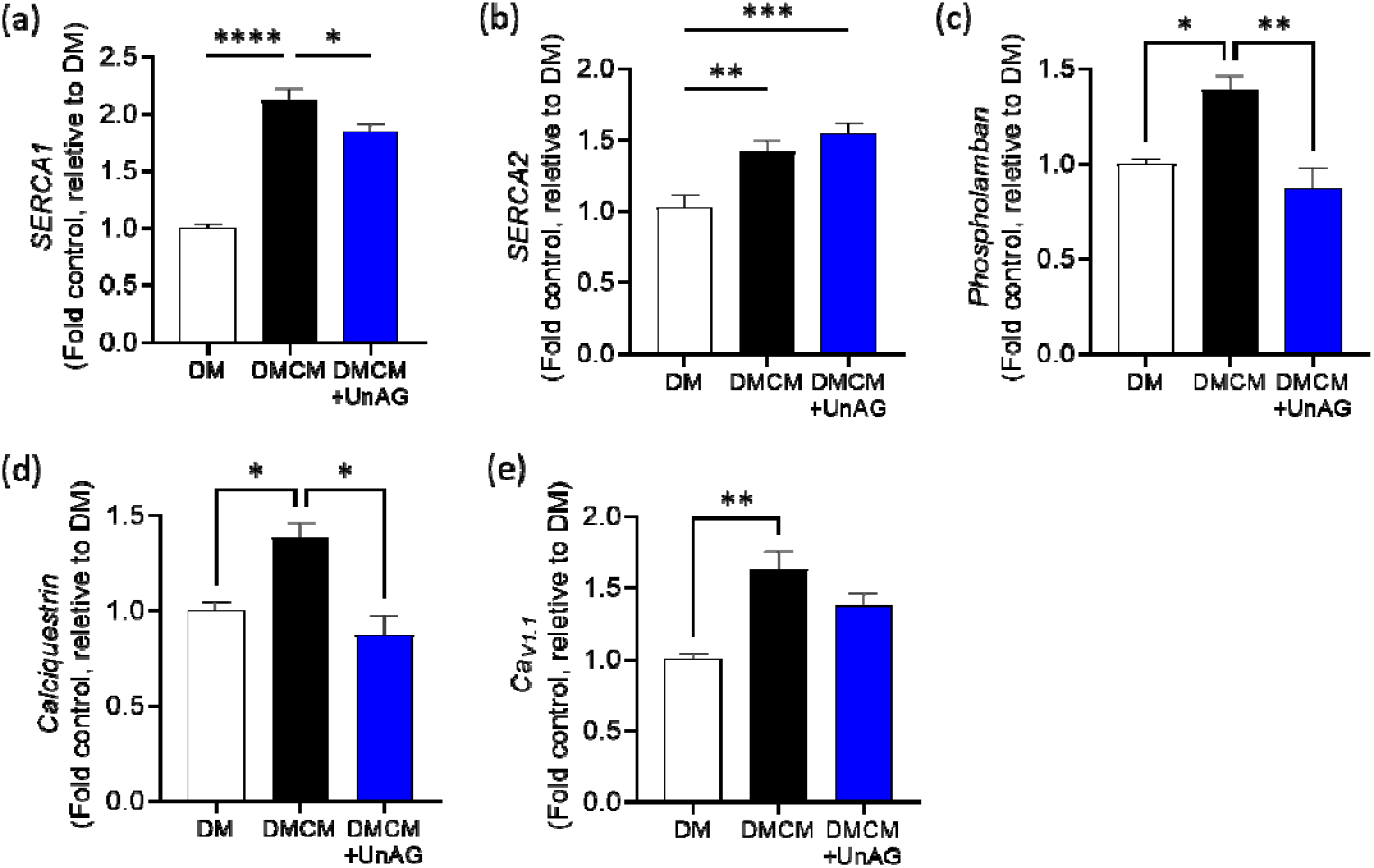
mRNA expression of cultured myotubes incubated with UnAG. The mRNA expression measured for genes related to calcium regulation, including *SERCA1* (a), *SERCA2* (b), *phospholamban* (c), *calciquestrin* (d), and voltage-dependent calcium channel L-type, *C*_*AV1*.*1*_ (e). One-way ANOVA followed by Tukey post hoc tests. *p<0.05. **p<0.01. ***p<0.001. ****p<0.0001. Bars represent means ±SEM. Abbreviations: SERCA, sarco-endoplasmic reticulum calcium ATPase; Ca_v_1.1, calcium channel, voltage-dependent alpha 1S subunit.

**Supplemental Table 1.**
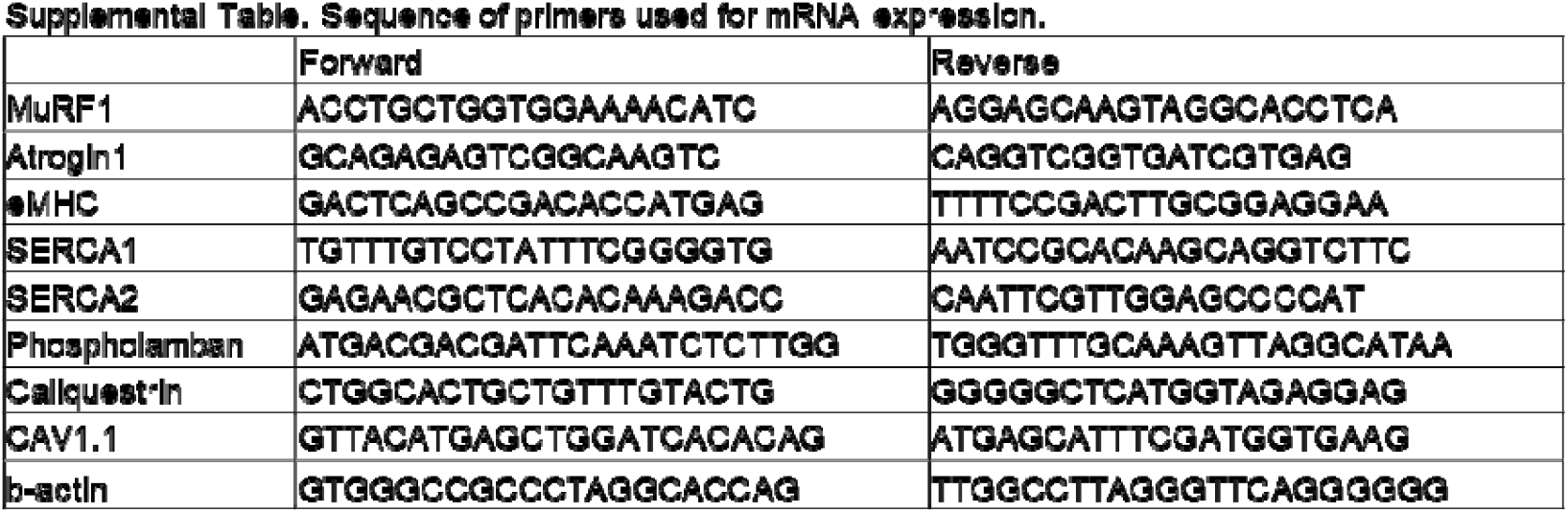
Primer sequences used for mRNA expressions based on literature.

## Notes

### Competing Interest Statement

The authors have declared no competing interest.

## Reference

1. Fearon K, Strasser F, Anker SD, Bosaeus I, Bruera E, Fainsinger RL, et al. Definition and classification of cancer cachexia: an international consensus. Lancet Oncol. 2011;12:489–95. doi:10.1016/S1470-2045(10)70218-7

2. Akezaki Y, Kikuuchi M, Hamada K, Ookura M. Incidence of cachexia in patients with advanced gastrointestinal cancer at the beginning of rehabilitation intervention. J Phys Ther Sci. 2020;32:16–9. doi:10.1589/jpts.32.16

3. Baracos VE, Martin L, Korc M, Guttridge DC, Fearon KCH. Cancer-associated cachexia. Nat Rev Dis Primers. 2018;4:doi:10.1038/nrdp.2017.105

4. Poulia KA, Sarantis P, Antoniadou D, Koustas E, Papadimitropoulou A, Papavassiliou AG, et al. Pancreatic Cancer and Cachexia-Metabolic Mechanisms and Novel Insights. Nutrients. 2020;12: doi:ARTN 154310.3390/nu12061543

5. Kojima M, Hosoda H, Date Y, Nakazato M, Matsuo H, Kangawa K. Ghrelin is a growth-hormone-releasing acylated peptide from stomach. Nature. 1999;402:656–60. doi:Doi 10.1038/45230

6. Takayama K, Katakami N, Yokoyama T, Atagi S, Yoshimori K, Kagamu H, et al. Anamorelin (ONO-7643) in Japanese patients with non-small cell lung cancer and cachexia: results of a randomized phase 2 trial. Support Care Cancer. 2016;24:3495–505. doi:10.1007/s00520-016-3144-z

7. Temel JS, Abernethy AP, Currow DC, Friend J, Duus EM, Yan Y, et al. Anamorelin in patients with non-small-cell lung cancer and cachexia (ROMANA 1 and ROMANA 2): results from two randomised, double-blind, phase 3 trials. Lancet Oncol. 2016;17:519–31. doi:10.1016/S1470-2045(15)00558-6

8. Katakami N, Uchino J, Yokoyama T, Naito T, Kondo M, Yamada K, et al. Anamorelin (ONO-7643) for the Treatment of Patients With Non-Small Cell Lung Cancer and Cachexia: Results From a Randomized, Double-Blind, Placebo-Controlled, Multicenter Study of Japanese Patients (ONO-7643-04). Cancer-Am Cancer Soc. 2018;124:606–16. doi:10.1002/cncr.31128

9. Ranjit R, Van Remmen H, Ahn B. Acylated Ghrelin Receptor Agonist HM01 Decreases Lean Body and Muscle Mass, but Unacylated Ghrelin Protects against Redox-Dependent Sarcopenia. Antioxidants (Basel). 2022;11:doi:10.3390/antiox11122358

10. Kim H, Ranjit R, Claflin DR, Georgescu C, Wren JD, Brooks SV, et al. Unacylated Ghrelin Protects Against Age-Related Loss of Muscle Mass and Contractile Dysfunction in Skeletal Muscle. Aging Cell. 2024;e14323. doi:10.1111/acel.14323

11. Rossetti A, Togliatto G, Rolo AP, Teodoro JS, Granata R, Ghigo E, et al. Unacylated ghrelin prevents mitochondrial dysfunction in a model of ischemia/reperfusion liver injury. Cell Death Discov. 2017;3:17077. doi:10.1038/cddiscovery.2017.77

12. Sheriff S, Kadeer N, Joshi R, Friend LA, James JH, Balasubramaniam A. Des-acyl ghrelin exhibits pro-anabolic and anti-catabolic effects on C2C12 myotubes exposed to cytokines and reduces burn-induced muscle proteolysis in rats. Mol Cell Endocrinol. 2012;351:286–95. doi:10.1016/j.mce.2011.12.021

13. Cappellari GG, Semolic A, Ruozi G, Vinci P, Guarnieri G, Bortolotti F, et al. Unacylated ghrelin normalizes skeletal muscle oxidative stress and prevents muscle catabolism by enhancing tissue mitophagy in experimental chronic kidney disease. Faseb Journal. 2017;31:5159-+. doi:10.1096/fj.201700126R

14. Cappellari GG, Zanetti M, Semolic A, Vinci P, Ruozi G, Falcione A, et al. Unacylated Ghrelin Reduces Skeletal Muscle Reactive Oxygen Species Generation and Inflammation and Prevents High-Fat Diet-Induced Hyperglycemia and Whole-Body Insulin Resistance in Rodents. Diabetes. 2016;65:874–86. doi:10.2337/db15-1019

15. Porporato PE, Filigheddu N, Reano S, Ferrara M, Angelino E, Gnocchi VF, et al. Acylated and unacylated ghrelin impair skeletal muscle atrophy in mice. J Clin Invest. 2013;123:611–22. doi:10.1172/Jci39920

16. Filigheddu N, Gnocchi VF, Coscia M, Cappelli M, Porporato PE, Taulli R, et al. Ghrelin and des-acyl ghrelin promote differentiation and fusion of C2C12 skeletal muscle cells. Mol Biol Cell. 2007;18:986–94. doi:10.1091/mbc.E06-05-0402

17. Lear PV, Iglesias MJ, Feijóo-Bandín S, Rodríguez-Penas D, Mosquera-Leal A, García-Rúa V, et al. Des-Acyl Ghrelin Has Specific Binding Sites and Different Metabolic Effects from Ghrelin in Cardiomyocytes. Endocrinology. 2010;151:3286–98. doi:10.1210/en.2009-1205

18. Kim H, Ranjit R, Claflin DR, Georgescu C, Wren JD, Brooks SV, et al. Unacylated Ghrelin Protects Against Age-Related Loss of Muscle Mass and Contractile Dysfunction in Skeletal Muscle. Aging Cell. 2024;23:doi:10.1111/acel.14323

19. Ahn B, Ranjit R, Premkumar P, Pharaoh G, Piekarz KM, Matsuzaki S, et al. Mitochondrial oxidative stress impairs contractile function but paradoxically increases muscle mass via fibre branching. J Cachexia Sarcopenia Muscle. 2019;10:411–28. doi:10.1002/jcsm.12375

20. Ahn B, Beharry AW, Frye GS, Judge AR, Ferreira LF. NAD(P)H oxidase subunit p47phox is elevated, and p47phox knockout prevents diaphragm contractile dysfunction in heart failure. Am J Physiol Lung Cell Mol Physiol. 2015;309:L497–505. doi:10.1152/ajplung.00176.2015

21. Roberts BM, Frye GS, Ahn B, Ferreira LF, Judge AR. Cancer cachexia decreases specific force and accelerates fatigue in limb muscle. Biochem Bioph Res Co. 2013;435:488–92. doi:10.1016/j.bbrc.2013.05.018

22. Ahn B, Ranjit R, Kneis P, Xu HY, Piekarz KM, Freeman WM, et al. Scavenging mitochondrial hydrogen peroxide by peroxiredoxin 3 overexpression attenuates contractile dysfunction and muscle atrophy in a murine model of accelerated sarcopenia. Aging Cell. 2022;21:doi:ARTN e1356910.1111/acel.13569

23. Ahn B, Pharaoh G, Premkumar P, Huseman K, Ranjit R, Kinter M, et al. Nrf2 deficiency exacerbates age-related contractile dysfunction and loss of skeletal muscle mass. Redox Biology. 2018;17:47–58. doi:10.1016/j.redox.2018.04.004

24. Herbst A, Widjaja K, Nguy B, Lushaj EB, Moore TM, Hevener AL, et al. Digital PCR Quantitation of Muscle Mitochondrial DNA: Age, Fiber Type, and Mutation-Induced Changes. J Gerontol A Biol Sci Med Sci. 2017;72:1327–33. doi:10.1093/gerona/glx058

25. Roberts BM, Ahn B, Smuder AJ, Al-Rajhi M, Gill LC, Beharry AW, et al. Diaphragm and ventilatory dysfunction during cancer cachexia. Faseb Journal. 2013;27:2600–10. doi:10.1096/fj.12-222844

26. Dasgupta A, Shukla SK, Vernucci E, King RJ, Abrego J, Mulder SE, et al. SIRT1-NOX4 signaling axis regulates cancer cachexia. Journal of Experimental Medicine. 2020;217:doi:ARTN e2019074510.1084/jem.20190745

27. Cosper PF, Leinwand LA. Myosin heavy chain is not selectively decreased in murine cancer cachexia. Int J Cancer. 2012;130:2722–7. doi:10.1002/ijc.26298

28. de Castro GS, Simoes E, Lima JDCC, Ortiz-Silva M, Festuccia WT, Tokeshi F, et al. Human Cachexia Induces Changes in Mitochondria, Autophagy and Apoptosis in the Skeletal Muscle. Cancers. 2019;11:doi:ARTN 126410.3390/cancers11091264

29. Brown JL, Lawrence MM, Ahn B, Kneis P, Piekarz KM, Qaisar R, et al. Cancer cachexia in a mouse model of oxidative stress. J Cachexia Sarcopenia Muscle. 2020;11:1688–704. doi:10.1002/jcsm.1261530.

30. Pharaoh G, Sataranatarajan K, Street K, Hill S, Gregston J, Ahn B, et al. Metabolic and Stress Response Changes Precede Disease Onset in the Spinal Cord of Mutant SOD1 ALS Mice. Front Neurosci. 2019;13:487. doi:10.3389/fnins.2019.00487

31. Song T, McNamara JW, Ma W, Landim-Vieira M, Lee KH, Martin LA, et al. Fast skeletal myosin-binding protein-C regulates fast skeletal muscle contraction. Proc Natl Acad Sci U S A. 2021;118:doi:10.1073/pnas.2003596118

32. Baracos VE, Reiman T, Mourtzakis M, Gioulbasanis I, Antoun S. Body composition in patients with non-small cell lung cancer: a contemporary view of cancer cachexia with the use of computed tomography image analysis. Am J Clin Nutr. 2010;91:1133S–7S. doi:10.3945/ajcn.2010.28608C

33. Xu H, Ahn B, Van Remmen H. Impact of aging and oxidative stress on specific components of excitation contraction coupling in regulating force generation. Sci Adv. 2022;8:eadd7377. doi:10.1126/sciadv.add7377

34. Kunz HE, Port JD, Kaufman KR, Jatoi A, Hart CR, Gries KJ, et al. Skeletal muscle mitochondrial dysfunction and muscle and whole body functional deficits in cancer patients with weight loss. J Appl Physiol. 2022;132:388–401. doi:10.1152/japplphysiol.00746.2021

35. Huot JR, Baumfalk D, Resendiz A, Bonetto A, Smuder AJ, Penna F. Targeting Mitochondria and Oxidative Stress in Cancer- and Chemotherapy-Induced Muscle Wasting. Antioxid Redox Signal. 2023;38:352–70. doi:10.1089/ars.2022.0149

36. Pin F, Huot JR, Bonetto A. The Mitochondria-Targeting Agent MitoQ Improves Muscle Atrophy, Weakness and Oxidative Metabolism in C26 Tumor-Bearing Mice. Front Cell Dev Biol. 2022;10:861622. doi:10.3389/fcell.2022.861622

37. Argilés JM, López-Soriano FJ, Stemmler B, Busquets S. Cancer-associated cachexia - understanding the tumour macroenvironment and microenvironment to improve management. Nat Rev Clin Oncol. 2023;20:250–64. doi:10.1038/s41571-023-00734-5

38. Herbst A, Prior SJ, Lee CC, Aiken JM, McKenzie D, Hoang A, et al. Skeletal muscle mitochondrial DNA copy number and mitochondrial DNA deletion mutation frequency as predictors of physical performance in older men and women. Geroscience. 2021;43:1253–64. doi:10.1007/s11357-021-00351-z

39. Larsen S, Nielsen J, Hansen CN, Nielsen LB, Wibrand F, Stride N, et al. Biomarkers of mitochondrial content in skeletal muscle of healthy young human subjects. J Physiol. 2012;590:3349–60. doi:10.1113/jphysiol.2012.230185

40. Ericson NG, Kulawiec M, Vermulst M, Sheahan K, O’Sullivan J, Salk JJ, et al. Decreased mitochondrial DNA mutagenesis in human colorectal cancer. PLoS Genet. 2012;8:e1002689. doi:10.1371/journal.pgen.1002689

